# Oxytocin treats respiratory depression and reduces mortality from fentanyl and the combination of xylazine-fentanyl

**DOI:** 10.64898/2026.02.23.702914

**Authors:** Joan B. Escobar, John Wainwright, Xin Wang, Olga Dergacheva, Matthew W. Kay, John R. Bethea, Vivek Jain, Vsevolod Y. Polotsky, David Mendelowitz

## Abstract

Opioid addiction and misuse are a serious national crisis that affects public health, as well as social and economic welfare. Mortality due to opioid misuse is further exasperated by the combination of opioids with non-opioid respiratory depressants such as xylazine that are resistant to mu opioid receptor antagonists such as naloxone. This study tested the hypothesis that oxytocin can mitigate the severe opioid induced respiratory depression (OIRD) and mortality induced by high doses of fentanyl or the combination of fentanyl with xylazine. Our results show OXT can improve survival and respiratory function in both male and female rats with opioid induced respiratory depression caused by fentanyl, as well as a combination of fentanyl and xylazine. The improvement in respiratory function by OXT post fentanyl-xylazine was significantly greater than the recovery using only naloxone. Chemogenetic activation of OXT receptor positive neurons in the ventral respiratory group (VRG) provided similar benefits to that of OXT administration in reversing OIRD. These results indicate OXT is a promising therapeutic target for reversing OIRD and the respiratory depression that occurs with the combination of opioids and xylazine, a situation where naloxone is only partially effective. Additional translational benefits of OXT include it can be repurposed as it is already a FDA approved drug for other uses, has a high safety profile, and is unlikely to induce the withdrawal or reversal of analgesia that occurs with naloxone.

**Key Points:**

- Oxytocin (OXT) improves survival and respiratory function in both male and female rats with opioid induced respiratory depression (OIRD) caused by fentanyl
- OXT also reverses OIRD induced by the combination of fentanyl and xylazine
- The improvement in respiratory function by OXT post fentanyl-xylazine was significantly greater than the recovery using only naloxone
- Chemogenetic activation of OXT receptor positive neurons in the ventral respiratory group (VRG) provided similar benefits to that of OXT administration in reversing OIRD
- These results indicate OXT is a promising therapeutic target for reversing OIRD and the respiratory depression that occurs with the combination of opioids and xylazine

## Introduction

Opioid addiction and misuse are a serious national crisis that affects public health, as well as social and economic welfare. The opioid crisis has been greatly exacerbated by the increased availability of synthetic opioids, such as fentanyl, and by the increased prescribing of opioid pain relievers (Morone and Weiner, 2013; Van Zee, 2009). Opioid overdose kills 130 people in the United States every day according to the CDC/NCHS National Vital Statistics System. The primary cause of death associated with opioids is opioid-induced respiratory depression (OIRD).

Naloxone, a competitive antagonist of mu-opioid receptors (MORs), has a rapid onset and helps reverse OIRD but is short acting and re-dosing is necessary to reverse long-acting synthetic opioids (Britch and Walsh, 2022; Pasternak and Pan, 2013; Wong et al., 2017). In addition, the toxic effects of fentanyl and its analogues also can include compromised breathing due to mechanisms not reversible by MOR antagonists (Torralva and Janowsky, 2019). Naloxone blocks the analgesic effects of opioids (Orman and Keating, 2009a, b) and causes acute withdrawal that often causes agitation, vomiting and diarrhea, limiting its use (Handal et al., 1983; Weisshaar et al., 2020). Pulmonary edema has also been reported after high doses of naloxone (Conley et al., 2024; Joseph et al., 2024; Kummer et al., 2022). Another MOR antagonist, nalmefene, has longer half-life (8-11 h), but it has limitations and adverse effects similar to naloxone (Britch and Walsh, 2022).

In addition, fentanyl is also being increasingly combined with other non-opioid respiratory depressants. Xylazine, an alpha-2 adrenergic receptor agonist is typically used as a sedative and analgesic in veterinary medicine, is not a controlled substance and is naloxone-resistant (Friedman et al., 2022). A fentanyl-xylazine mixture (i.e., “tranq-dope”) represents a rapidly emerging public health threat and was present in 6.7% of opioid related overdose deaths in 2020. A nonlethal dose of xylazine (100 mg/kg) produces a synergistic interaction with fentanyl and decreased the estimated LD50 dose for fentanyl by approximately 100-fold (Smith et al., 2023). Alternative, non-opioid receptor antagonist-based approaches for opioid and other drug induced respiratory depression is needed.

Recent work has suggested that oxytocin can act as a respiratory stimulant in patients with obstructive sleep apnea (Jain et al., 2020; Jain et al., 2017). This study tested the hypothesis that oxytocin can mitigate severe OIRD and mortality in unrestrained conscious animals induced by high doses of fentanyl or the combination of fentanyl with xylazine. This work also examined if there are any sex differences in OIRD caused by fentanyl and treatment with OXT. To identify possible sites of action of oxytocin we also tested if chemogenetic activation of oxytocin receptor positive neurons in the ventral respiratory group (VRG) could reverse OIRD.

## Methods

### Animals

All animal experiments were performed in accordance with NIH and Institutional Animal Care and Use Guidelines approved by the George Washington University Institutional Animal Care and Use Committee (IACUC, protocols #2022-028, #2022-035, #2025-141). All animals were housed in standard environmental conditions (24°C–26°C in the 12-h light/dark cycle, 7a.m.–7p.m. lights on), water and food were available ad libitum. Unrestrained Whole-Body Plethysmography (Scireq, Montreal, Canada) recordings were conducted to quantify respiration. Following a 30-minute acclimation period, animals received an I.P. injection of fentanyl or the combination of fentanyl and xylazine followed 10-15 minutes later by an injection of either saline, Oxytocin (200 nmol/kg), Naloxone (0.1 mg/kg), the combination of oxytocin and naloxone. or the DREADDS agonist Clozapine N-Oxide (1 mg/kg). Respiratory function was reassessed one hour after the last injection.

For chemogenetic activation of OXTR+ neurons in the VRG the transgenic OXT receptor (OXTR)-Cre mouse line (B6.Cg-Oxtrtm1.1(cre)Hze/J, Common Name: Oxtr-T2A-Cre-D, Jackson Labs stock # 031303) was used. An AAV floxed excitatory DREADDs vector (AAV8-hSyn-DIO-hM3D(Gq)-mCherry, ≥5 x 1012 171 vg/mL, UNC Core) was injected into the VRG to selectively express excitatory DREADDs in OXTR+ VRG neurons in male and female mice. For this injection mice were anesthetized using a mixture of ketamine (100 mg/kg, i.p.) and xylazine (10 mg/kg, i.p.), then secured in a stereotaxic frame with the neck bent at 45 degrees to expose the calamus scriptorius (David Kopf Instruments, CA, USA). Using stereotaxic guidance of a pulled glass capillary (40μm tip diameter) (G1, Narishige, UK) attached to a pneumatic microinjector (IM-11-2, Narishige, UK), 200nL of AAV floxed DREADDs was bilaterally injected into the VRG (AP 0.4mm from the Obex, ML 1.5mm, DV 4.5mm). The capillary remained in place for 5 minutes to allow diffusion and was then removed slowly to avoid dispersion to neighboring brainstem regions. DREADDs-mCherry expression was observed in the VRG in all mice used for these experiments after the mice were sacrificed.

#### Brain slice preparation, Immunohistochemistry, and confocal images

After completion of Whole-Body Plethysmography recordings, animals were anesthetized by isoflurane and transcardially perfused with phosphate buffered saline (PBS) followed by 4% paraformaldehyde (PFA). Brains were dissected, post-fixed in 4% PFA overnight at room temperature, then washed 3x10 min in PBS. A series of 50 μm coronal section medullary slices containing the VRG were obtained using a Leica dissection vibratome (Leica VT 1000S). 10-20 VRG medullary slices were obtained from each animal. All slice sections were processed using immunohistochemistry for mCherry and Phox2b to identify Cre-dependent VRG and phox2b positive-expressing neurons, respectively. After blocking nonspecific proteins in 10% normal goat serum (NGS) in PBS with 0.3% Triton X-100 (PBST) for 4 hours at room temperature, slices were incubated in primary antibody at 4°C for 24 hours (Chicken anti-RFP antibody (1:1000, Millipore Sigma, Cat. # AB3528), and Mouse monoclonal anti Phox2b (1:200, sc-376997; Santa Cruz Biotechnology, Inc., Santa Cruz, CA)). Secondary antibodies were applied for 4 hours at room temperature. Secondary antibodies were Goat anti Chicken Alexa Fluor™ Plus 555, (Invitrogen, Cat. # A32932) and goat anti mouse Alexa Fluor 647 (Invitrogen, Cat. # A-21241), both 1:200 dilution. Slides were mounted with Fluorogel (Electron Microscopy Sciences, SKU: 17985-10) and imaged with a Leica TCS SP8 multi-photon confocal microscope equipped with supercontinuum white laser source and single molecule detection hybrid detectors (SMD HyD, Leica, Wetzlar, Germany). Leica TCS SP8 MP and Zeiss LSM 980 confocal microscopy were used to assess colocalization of mCherry and Phox2b labeled neurons in the VRG. Brain tissue slices containing VRG were examined with 555 and 647 nm wavelengths to visualize mCherry-Alexa Fluor 555 and the Phox2b-Alexa Fluor 647, respectively. Images were captured with a DFC365FX camera at 2048 by 2048-pixel resolution. Full field images of the entire slice were taken at 10x to localize the VRG. Z-stacks were then taken with 20x/0.75 oil-immersion objective, at z-step size of 0.9 μm to produce image volumes allowing for the identification of colocalization of mCherry and Phox2b positive neurons. Images were processed and analyzed using Imaris 10 software (Oxford Instruments Oxford, UK).

### Data Analysis

All data are presented as mean ± SEM. For *ex vivo* experiments, ‘n’ is reported as number of identified neurons in the VRG. For *in vivo* experiments, ‘n’ is the number of animals. These values are stated throughout the Results and Figure Legends. Statistical comparisons were made using repeated-measurements (RM) one-way and two-way ANOVA with Dunnett’s and Tukey’s multiple comparisons, paired or unpaired Student’s tests, as appropriate. Specific statistical tests are noted in the results. Differences were considered statistically significant if p-value *<* 0.05. Plethysmography data was obtained using the Emka IOX software (version 2.10.5.28, Emka Technologies, Sterling VA). Graph creation and statistical analyses were conducted using GraphPad Prism 9 (GraphPad Software, San Diego, CA).

## Results

Injection of fentanyl (0.5 mg/kg) in male and female rats induced rapid respiratory depression with decreases in respiratory rate (figure 1, top) as well as an increased incidence of apneas (figure 1, bottom). Injection of oxytocin (OXT, 200 mmol/kg, ip) significantly improved respiratory rate and decreased the incidence of apneas (figure 1). There were no significant differences between males (figure 1 left) and females (figure 1, right) in either the responses to fentanyl or beneficial actions of OXT.

**Figure 1.**
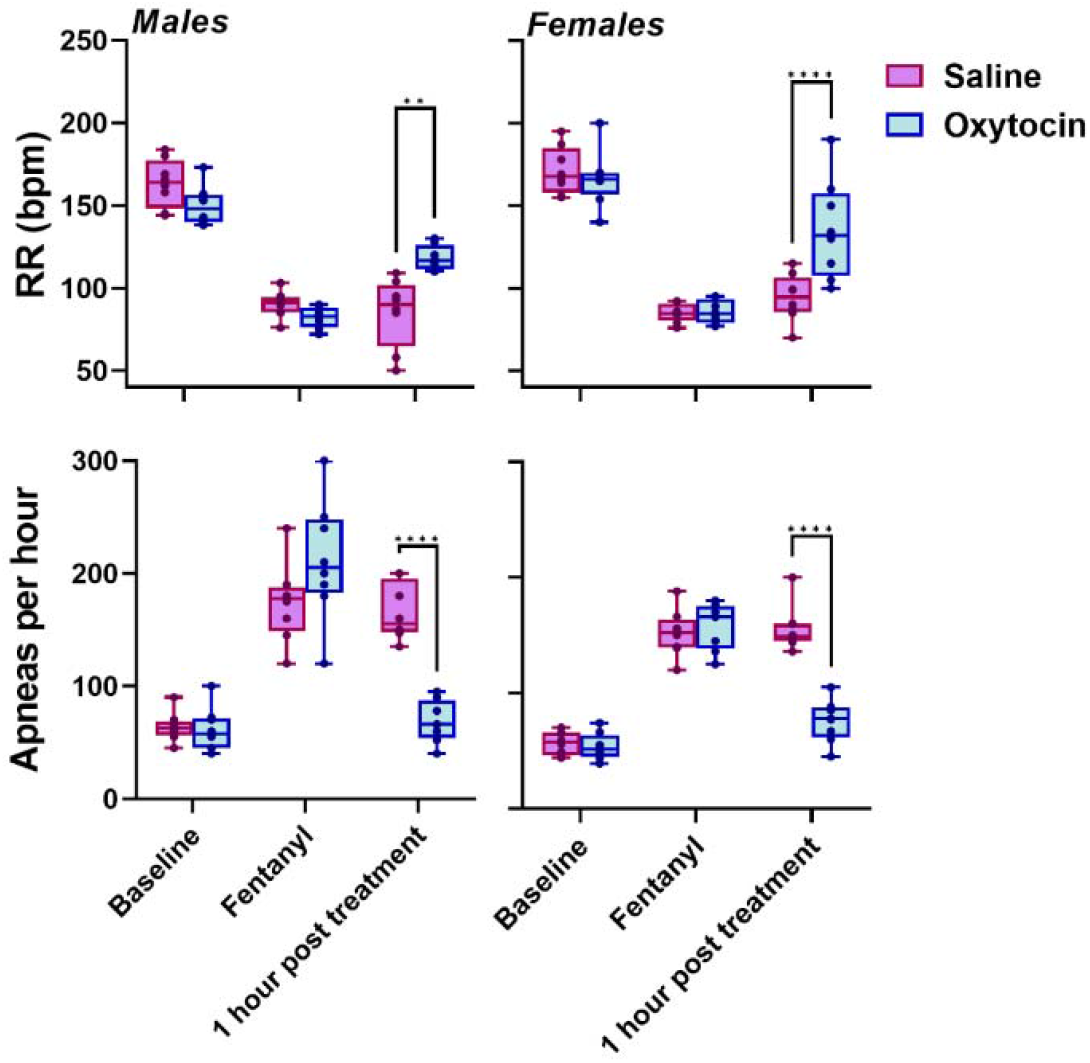
Effect of intraperitoneal (IP) oxytocin (200 nmol/kg) on fentanyl-induced respiratory depression. Respiratory rate (upper panels) and apnea occurrence (lower panels) are shown for males (left) and females (right) at baseline, after IP fentanyl injection (0.5 mg/kg), and 1 hour after treatment with oxytocin or saline. Statistical significance is indicated by asterisks for post hoc comparisons between the saline and oxytocin groups at 1 hour post-treatment. *n* = 8 per group.

Using a high dose of fentanyl (1.3 mg/kg) respiratory depression was more severe and without treatment the survival at 5 hours post fentanyl administration in male rats was 55%, whereas with OXT treatment there was 100% survival (figure 2, left). Female rats had a significantly higher rate of survival than males without treatment at this high dose (1.3 mg/kg), and, similar to males, with OXT treatment there was 100% survival in female rats (figure 2, right).

**Figure 2.**
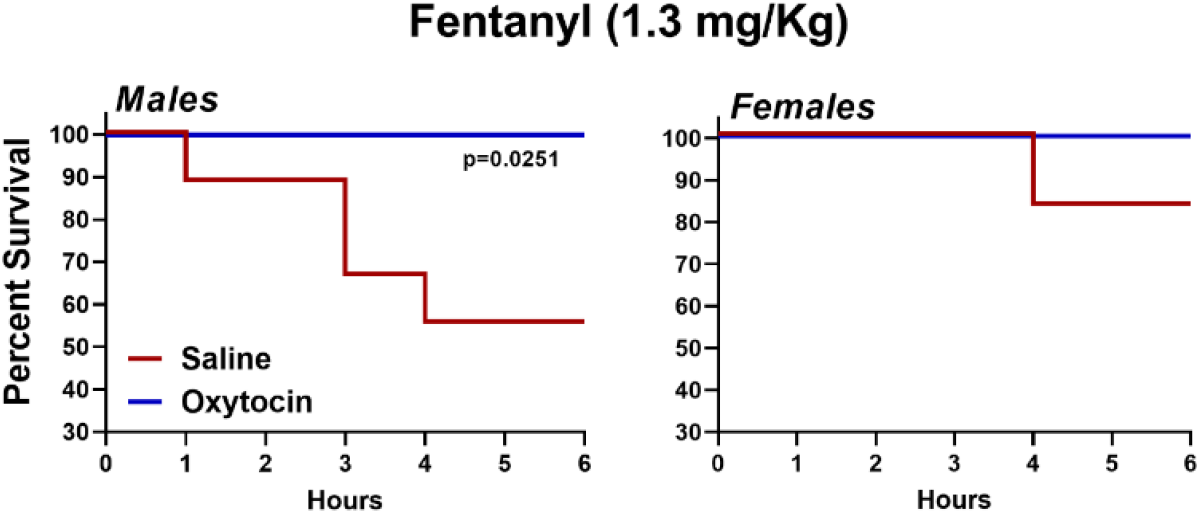
Mortality following IP fentanyl injection (1.3 mg/kg) and subsequent treatment with oxytocin (200 nmol/kg) or saline in males (left) and females (right). Survival curves over 6 hours were compared using the Mantel–Cox (log-rank) test. *p* values are indicated where statistically significant differences were detected. *n* = 9 per group.

Administration of a combination of fentanyl and xylazine (0.5 mg/kg and 0.1 mg/kg, respectively) induced a more severe respiratory depression of respiratory rate and incidence of apneas compared to fentanyl by itself (figure 3).

**Figure 3.**
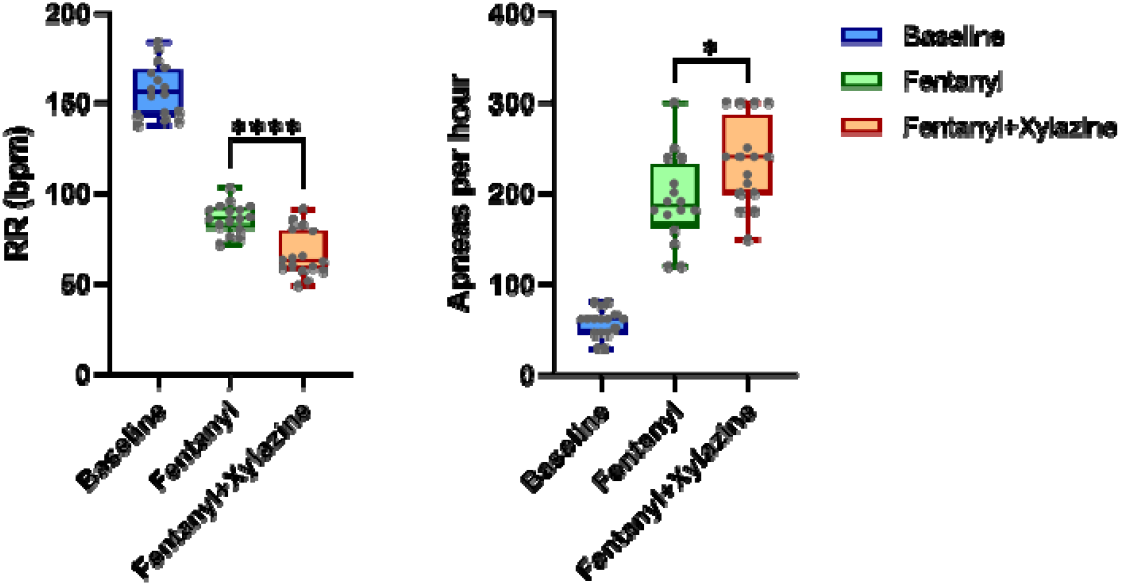
Respiratory depression induced by IP fentanyl (0.5 mg/kg) alone or fentanyl combined with xylazine (0.5 and 0.1 mg/kg, respectively). Respiratory rate (left) and apnea occurrence (right) are shown at baseline and after injection of fentanyl or fentanyl + xylazine. Statistical significance of post hoc comparisons between the fentanyl and fentanyl + xylazine groups is indicated by asterisks. *n* = 8 per group.

To compare the treatment of OXT with naloxone animals were given the combination of fentanyl and xylazine ((0.5 mg/kg and 0.1 mg/kg, respectively) followed by either naloxone (0.1 mg/kg, ip), OXT (200 mmol/kg, ip) or the combination of naloxone with OXT. As expected, both naloxone and OXT improved respiratory rate and reduced the incidence of apneas (figure 4), however suprisingly the improvement seen with OXT was significantly greater than that with naloxone. The treatment of OIRD caused by fentanyl and xylazine with the combination of naloxone and OXT was not significantly greater than OXT by itself (figure 4). The mortality caused by the combination of fentanyl and xylazine (1.3 mg/kg and 10 mg/kg, respectively) was prevented by either naloxone or OXT (figure 5).

**Figure 4.**
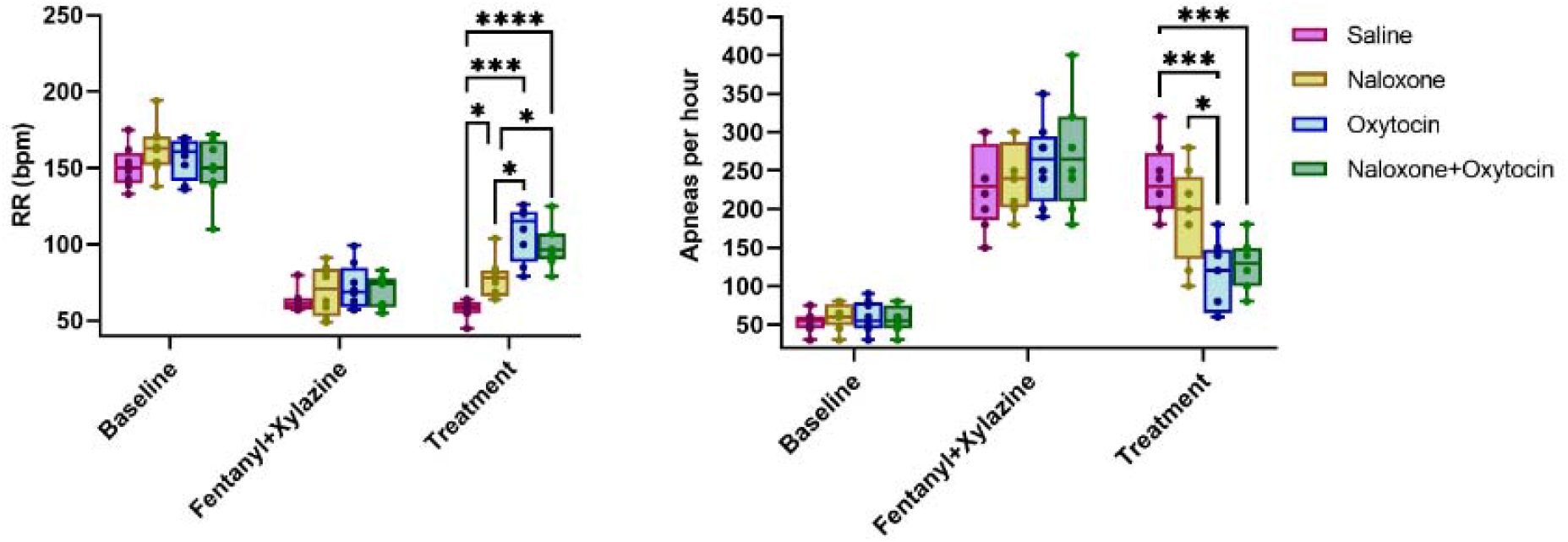
Effects of IP naloxone (0.1 mg/kg), oxytocin (200 nmol/kg), or their combination on respiratory depression induced by IP fentanyl + xylazine (0.5 and 0.1 mg/kg, respectively). Respiratory rate (left) and apnea occurrence (right) are shown at baseline, after fentanyl + xylazine injection, and 1 hour after treatment. Statistical significance is indicated by asterisks for multiple post hoc comparisons among treatment groups at 1 hour post-injection. *n* = 8 per group.

**Figure 5.**
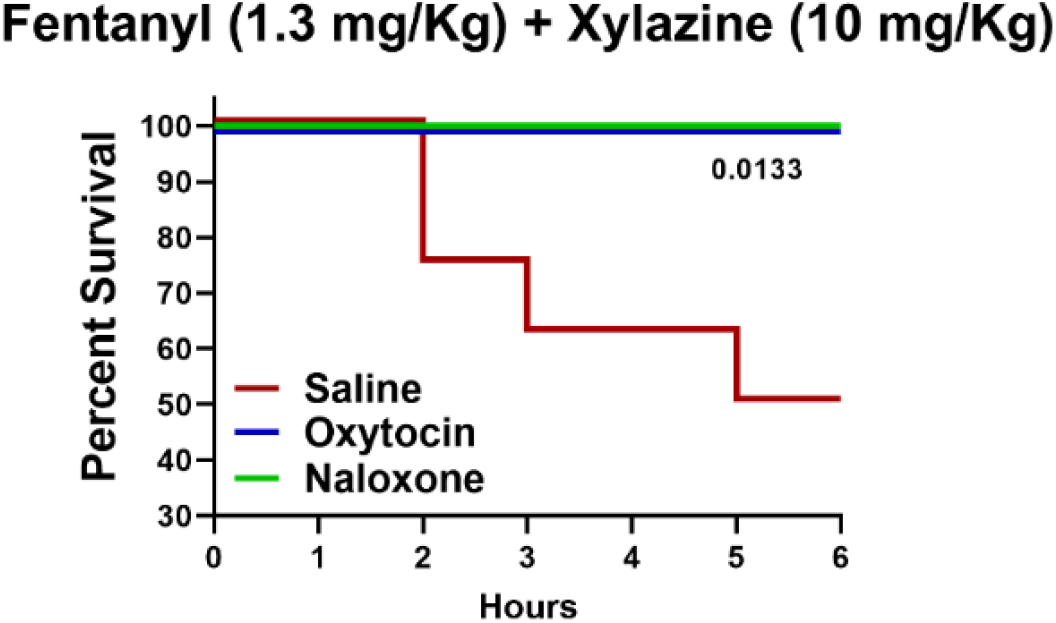
Mortality following IP fentanyl + xylazine injection (1.3 and 0.1 mg/kg, respectively) and subsequent treatment with oxytocin (200 nmol/kg), naloxone (0.1 mg/kg), or saline. Survival curves over 6 hours were compared using the Mantel–Cox (log-rank) test. *p* values are indicated where statistically significant differences were detected. *n* = 8 per group.

To identify possible sites of action of oxytocin we tested if chemogenetic activation of oxytocin receptor positive neurons in the VRG could reduce OIRD. In mice in which floxed excitatory DREADDs was injected into the VRG in OXTR+cre mice (figure 6A) administration of the DREADDs agonist CNO (1 mg/kg) significantly reversed the respiratory depression induced by fentanyl (10 mg/kg), see figure 6B. Confocal image analyses show only 4.3 percent of OXTR+ mCherry positive neurons in the VRG (figure 6C, representative OXTR+/mCherry+ expression seen in a total of 771 VRG neurons from 27 slices (5-7 slices/animal from 5 animals in total)) overlap with Phox2b positive cells that were more densely located medial to the OXTR+ neurons (figure 6C).

**Figure 6.**
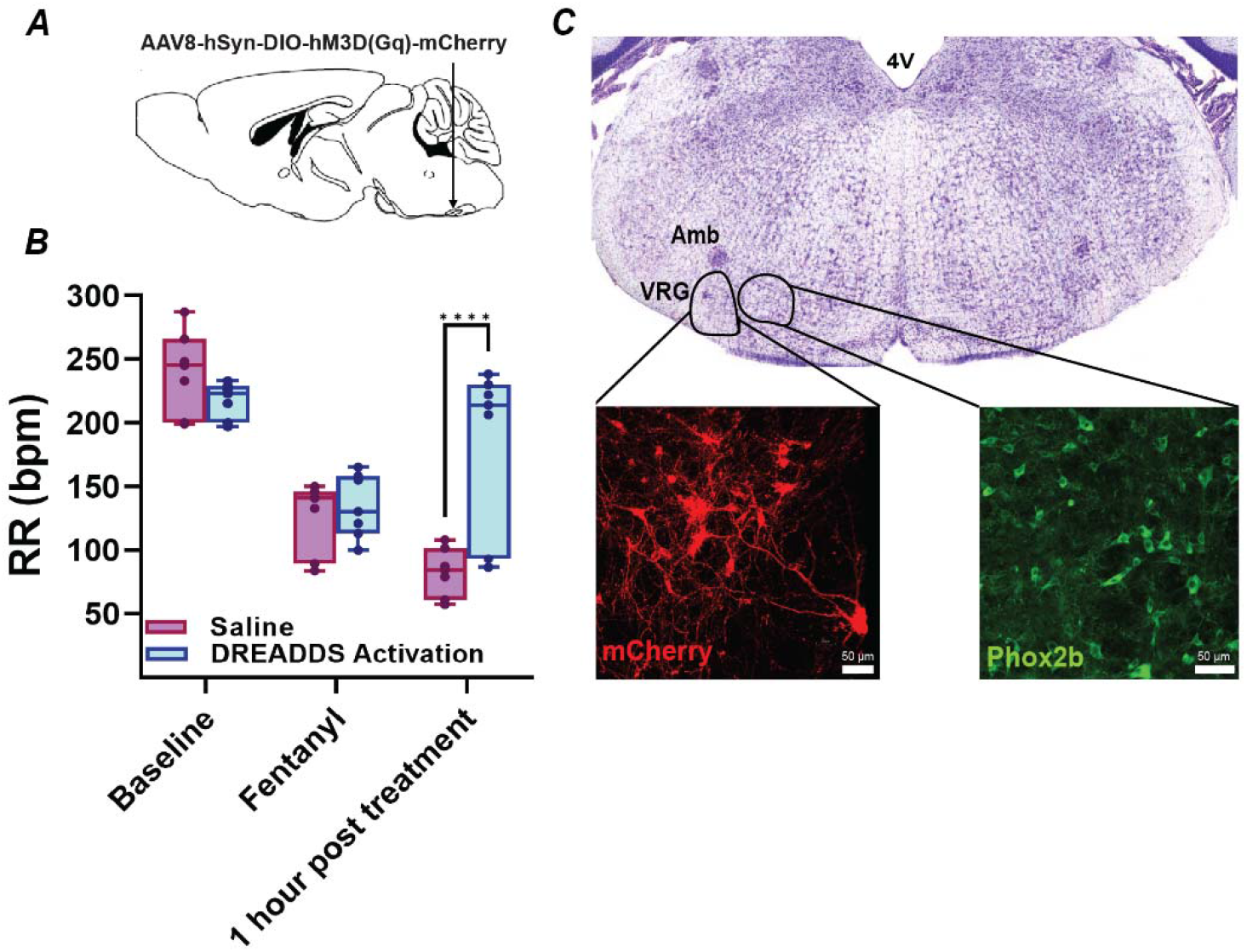
Effect of DREADDs activation on fentanyl-induced respiratory depression. OXTR+cre mice were injected with floxed excitatory DREADDs into the ventral respiratory group (A). The respiratory depression evoked by fentanyl (10 mg/kg) was reversed by IP clozapine-N-oxide (CNO, 1 mg/kg) to activate excitatory DREADDs in OXTR+ neurons. Respiratory rate is shown at baseline, after fentanyl injection, and 1 hour after CNO administration. Statistical significance is indicated by asterisks for post hoc comparisons between saline and DREADDs activation at 1-hour post-treatment. *n* = 7 per group. Confocal image analyses show only 4.3 percent of OXTR+ mCherry positive neurons in the VRG (figure 6C, representative OXTR+/mCherry+ expression seen in a total of 771 VRG neurons from 27 slices (5-7 slices/animal from 5 animals in total)) overlap with Phox2b positive cells that were more densely located medial to the OXTR+ neurons (figure 6C).

## Discussion

This study shows OXT can improve survival and respiratory function in animals with opioid induced respiratory depression caused by fentanyl, as well as a combination of fentanyl and xylazine. The improvement in respiratory function by OXT post fentanyl-xylazine was significantly greater than the recovery using naloxone. Chemogenetic activation of OXT receptor positive neurons in the VRG provided similar benefits to that of OXT administration in reversing OIRD. While the benefits of OXT was similar in both males and females, we found female rats had a higher survival after high doses of fentanyl compared to males. However unlike the results in this study, in which fentanyl was given via intraperitoneal injection, rats given a high intravenous dose (25 microg/kg) of fentanyl produced a respiratory depression that was slightly more severe in females, suggesting the route of administration might also play a role in sex differences (Marchette et al., 2023).

The findings in this study extends prior work that has shown OXT can stimulate respiration. OXT microinjected into the pre-Bötzinger complex within the VRG, a brainstem site postulated to be essential for generating inspiratory rhythm, increased respiratory frequency and diaphragm EMG activity (Kc and Dick, 2010). In addition, OXT may serve as a respiratory stimulant by activating neurons in the rostral ventrolateral medulla (RVLM) as microinjection of OXT into the RVLM also increased respiratory frequency and diaphragm muscle activity (Mack et al., 2002). Stimulating Phox2b+ retrotrapezoid nucleus (RTN) neurons, important in chemosensitivity, enhances breathing after fentanyl administration, whereas their inhibition exacerbates fentanyl induced hypoventilation (Moreira et al., 2025). Application of an oxytocin receptor agonist and optogenetic activation of oxytocinergic receptors/varicosities in the RTN stimulated respiration by increasing breathing amplitude (Araujo et al., 2025).

In addition to prior work that showed IV administration of oxytocin dose-dependently rescued fentanyl induced OIRD (Brackley and Toney, 2021) additional work has demonstrated intranasal (IN) OXT significantly increased the amplitude of inspiratory-related tongue muscle activity, and increased the firing of protruder hypoglossal motorneurons that would provide tongue protrusion and upper airway opening (Dergacheva et al., 2023). In clinical studies intranasal (IN) oxytocin reduces duration of apneas and hypopneas in patients with Obstructive Sleep Apnea (OSA) (Jain et al., 2020). The arterial oxygen desaturation that occurred during obstructive events was significantly (p<0.001) less severe and the risk of bradycardia associated with obstructive events was also significantly (p<0.002) reduced with IN OXT. In addition, IN OXT acted as a respiratory stimulant, increasing respiratory rate during non-obstructive periods (Jain et al., 2020).

In summary, these results indicate OXT is a promising therapeutic target for reversing opioid induced respiratory depression, as well as reversing the more severe respiratory depression that occurs with the combination of opioids and xylazine, a situation where naloxone is only partially effective. Additional translational benefits of OXT include it can be repurposed as it is already a FDA approved drug for other uses, has a high safety profile, and is unlikely to induce the withdrawal or the reversal of analgesia that occurs with using MOR antagonists.

## Acknowledgment

This work was supported by NIH grant UG3DA062506 and a NIH Shared Instrument Grant NIH S10OD032420

